# Microvascular plasticity in stroke recovery: Longitudinal snapshots, network statistical analysis, and dynamics

**DOI:** 10.1101/2023.06.29.547081

**Authors:** Samuel A Mihelic, Shaun A Engelmann, Mahdi Sadr, Chakameh Z Jafari, Annie Zhou, Michael R Williamson, Andrew K Dunn

## Abstract

This research article quantitatively investigates neuro-microvascular network remodeling dynamics following stroke using a novel in vivo two-photon angiography (cubic millimeter volume, weekly snapshots) and high throughput (thousands of connected capillaries) vascular vectorization method. The results suggest distinct temporal patterns of cerebrovascular plasticity, with acute remodeling peaking at one week post-stroke. The network architecture then gradually stabilizes, returning to a new steady state after four weeks. These findings align with previous literature on neuronal plasticity, highlighting the correlation between neuronal and neurovascular remodeling. Quantitative analysis of neurovascular networks using length- and strand-based statistical measures reveals intricate changes in network anatomy and topology. The distance and strand-length statistics show significant alterations, with a peak of plasticity observed at one week post-stroke, followed by a gradual return to baseline. The orientation statistic plasticity peaks at two weeks, gradually approaching the (conserved across subjects) stroke signature. The underlying mechanism of the vascular response (angiogenesis vs. tissue deformation), however, is yet unelucidated, requiring network registration advancements. Overall, the combination of two-photon angiography, vectorization, reconstruction/visualization, and statistical analysis enables both qualitative and quantitative assessments of neurovascular remodeling dynamics, demonstrating an impactful method for investigating neuro-microvascular network disorders and the therapeutic modes of action thereof. Understanding the timing and nature of neurovascular remodeling allows for optimized interventions, including personalized medicine for stroke rehabilitation. Additionally, the evaluation of pharmaceutical interventions using these tools may facilitate targeted drug development. Furthermore, neurovascular coupling dynamics have implications for neurodegenerative diseases, brain aging, and the field of brain-computer interfaces.

## 1 Introduction

The cerebrovasculature is a highly dense network of blood vessels that support metabolic activity in the brain through neurovascular coupling [25]. To briefly summarize, neuronal activity is accompanied by localized increases in blood flow brought forth by vasodilation events, enabling increased oxygen and nutrient exchange in active tissue. With this strong relationship between neuronal function and the cerebrovasculature, perturbation to the vascular structure can result in profound neurofunctional consequences. Due to neurovascular coupling, it is reasonable to hypothesize that neurovascular plasticity and neuronal plasticity would be highly correlated [42]. Therefore, it is paramount to thoroughly understand the intricacies of both healthy and unhealthy cerebrovascular structure and plasticity.

Endpoint histological studies have allowed identification of many conditions where structure deviates from normal. In humans, microvascular density declines as much as 10-30% in certain brain regions with age [8, 47, 36]. In mouse models with congenital deafness, a decrease in vascular branch density is identified in the auditory cortex while increases in density are present in both the visual and somatosensory barrel cortex [30]. For specimens that undergo chronic hypoxia, there is profound evidence for increases in vascular density [39, 6, 34]. Neurodegenerative diseases such as Alzheimer’s, Parkinson’s, and Huntington’s disease all also result in higher vessel density, whereas ALS results in a lower vessel density [7]. Molecular differences were detected between older and younger mouse populations to explain the effect of age on neurovascular plasticity in response to stroke recovery [4].

Endpoint studies have elucidated many important biological phenomena, however they naturally limit the temporal sampling to once per subject. Conclusions are therefore only drawn across specimens which obscures results and complicates findings. Additionally, the vascular structure after preservation does not perfectly represent the in vivo state which then requires careful calibration of microvessel diameter [28]. The issues of endpoint studies are addressed with chronic in vivo imaging making use of two-photon microscopy, but there are restrictions to the amount of vasculature that can be evaluated due to a limited lateral field of view (FOV) and imaging depth. Nevertheless, in vivo mouse studies have been performed in mice to investigate cerebrovascular plasticity in cases of aging [22], physical activity [17], hypoxia [34], and ischemia [53]. Conclusions from these mostly relied on manual evaluation of images which further decreased the already limited volumes that investigators were able to evaluate.

In vivo studies suggest that vascular structure in the brain is stable with a small but present level of microvessel turnover [17, 22]. However, in vivo imaging and analysis is technically difficult, relationships between stimuli and cerebrovascular plasticity are not always so clear, and studies require large sample sizes to establish statistical significance, which may be intractable without automated assistance. As an example, many endpoint studies in rodents reveal that maintained physical activity encourages at least temporary increases in microvascular density in the motor cortex and hippocampus [31, 50, 12, 52, 48]. However, no difference was identified between mice housed with monitored exercise wheels and a control group in a chronic in vivo evaluation performed by [17] using manual vessel tracing. Therefore it is important to repeat this study with an automated vectorization method to increase the statistical power.

Vectorization of microcirculatory angiograms enables three-dimensional reconstruction and quantitative comparison of samples, increasing the volumes of data and number of statistics that researchers can investigate. Ex vivo examples with high image quality are appropriate for image binarization follwed by skeletonization [9, 51, 5, 3]. These methods have been used to reconstruct an entire mouse brain [18], to enable blood flow simulations [49, 16], and to show organizational similarities in vascular structures across species [45].

In vivo neurovascular images have been vectorized previously by a few groups to study network topology [23, 35] in the context of stroke [21], simulate Laser Speckle Contrast Imaging (LSCI) acquisition [26, 27], and evaluate experimental imaging methods [58, 57]. However, this detail of analysis requires an imaging strategy that results in a sufficient signal to background ratio (SBR) and a vectorization strategy that is robust to local image quality fluctuations.

Here we demonstrate an in vivo imaging and vectorization strategy that covers large tissue volumes (exceeding 1 cubic millimeter) in the mouse brain at the resolution of capillaries (1.5 micrometers in *x*-*y*, 3 in *z*). Furthermore, we demonstrate a high-performance vectorization strategy that can accurately and efficiently compare the anatomy of the intricate microvascular network across samples in a longitudinal (five timepoints, once per week) stroke model study. To our knowledge, the level of detail of in vivo anatomical statistical analysis using automated vectorization techniques as presented here has been shown by [23, 21], however never before in the longitudinal and differential manner presented here. The imaging and vectorization methods enable a study of cerebral plasticity with high statistical significance and serve as an example for future animal model investigations into potential therapeutics to treat neurodegenerative diseases and neural injuries.

## 2 Materials and methods

### 2.1 Animal Preparation and Photothrombosis

Procedures were approved by the University of Texas Institutional Animal Care and Use Committee. Cranial windows were implanted on anesthetized (isoflurane, 1.5%, 0.6-0.8 liters per minute, LPM) male C57BL/6 mice while body temperature was maintained with a heating pad using a procedure similar to ones previously described [53, 40]. Carprofen was injected at 10mg/kg prior to surgery. A skull portion was then removed and a 5 mm diameter glass coverslip was implanted in its place using cyanoacrylate and dental cement following an injection of carprofen. Mice were housed 2-5 per cage both before and after the procedure. Mice were allowed to heal for three weeks prior to optical imaging and photothrombosis.

For ischemic mice, photothrombotic ischemia was induced through injecting 0.15ml of 15 mg/ml rose bengal retro-orbitally and irradiating a penetrating arteriole branching from the middle cerebral artery for 15 minutes using a 532 nm, 20mW laser source focused to a 300 um diameter spot size. Mice were anesthetized with isoflurane (1.5%, 0.6-0.8 LPM) and body temperature was maintained with a heating pad during photothrombosis. Pial anatomy was visualized using laser speckle contrast imaging during the procedure to select which artery to target and to confirm occlusion.

### 2.2 Laser Speckle Contrast Imaging

LSCI was performed in mice anesthetized with isoflurane (1.5%, 0.6-0.8 LPM) using a system described previously [46]. The brain surface was imaged onto a camera and illuminated using a 785 nm laser diode to provide light for LCSI. This imaging technique, which is sensitive to blood flow in the tissue, was primarily used to orient the brain prior to 2PM imaging sessions to encourage longitudinal reproducibility to image the same tissue volume.

### 2.3 2PM Optical Imaging

2PM imaging was performed immediately after LSCI, although we needed a different oxygen source for anesthetization (1.5%, 0,6-0.8 LPM), so slight differences in consciousness may be present. Texas red was injected retro-orbitally and excited using a previously described fiber amplifier. The microscope used had a resonant scanner to decrease acquisition time. Six stacks exceeding a 600 um depth, 3 um step size were acquired in adjacent, tiled regions and stitched together. Final images had approximately a 1.6×1.1×0.6mm FOV, resulting in a volume exceeding 1 mm3. Pial architecture was used to image the same region at each timepoint.

### 2.4 Experimental Design

The experimental methods and design are graphically explained in Figure 1. Laser Speckle Contrast Imaging (LSCI) was first performed approximately three weeks after cranial window implantation and used to evaluate craniotomy quality in order to select the most viable specimens for chronic study and stroke induction. All selected mice were then imaged again with both LSCI and two-photon microscopy (2PM) to get the initial anatomical and blood flow timepoints. The following day photothrombotic ischemia was induced through green light photothrombosis in a portion of the mice. Afterwards in these mice, weekly 2PM and LSCI was performed for four weeks resulting in 5 total timepoints (weeks 0-4). An additional LSCI time point was acquired 2 days after ischemia which helped quantify infarct size. In the healthy mice where no photothrombosis was performed, 2P and LSCI imaging were performed bi-weekly over the course of 4 weeks resulting in 3 timepoints (weeks zero, two, and four). Ultimately, images from one healthy control and three stroke model mice were vectorized and included in statistical (including directional) analysis.

**Fig 1.**
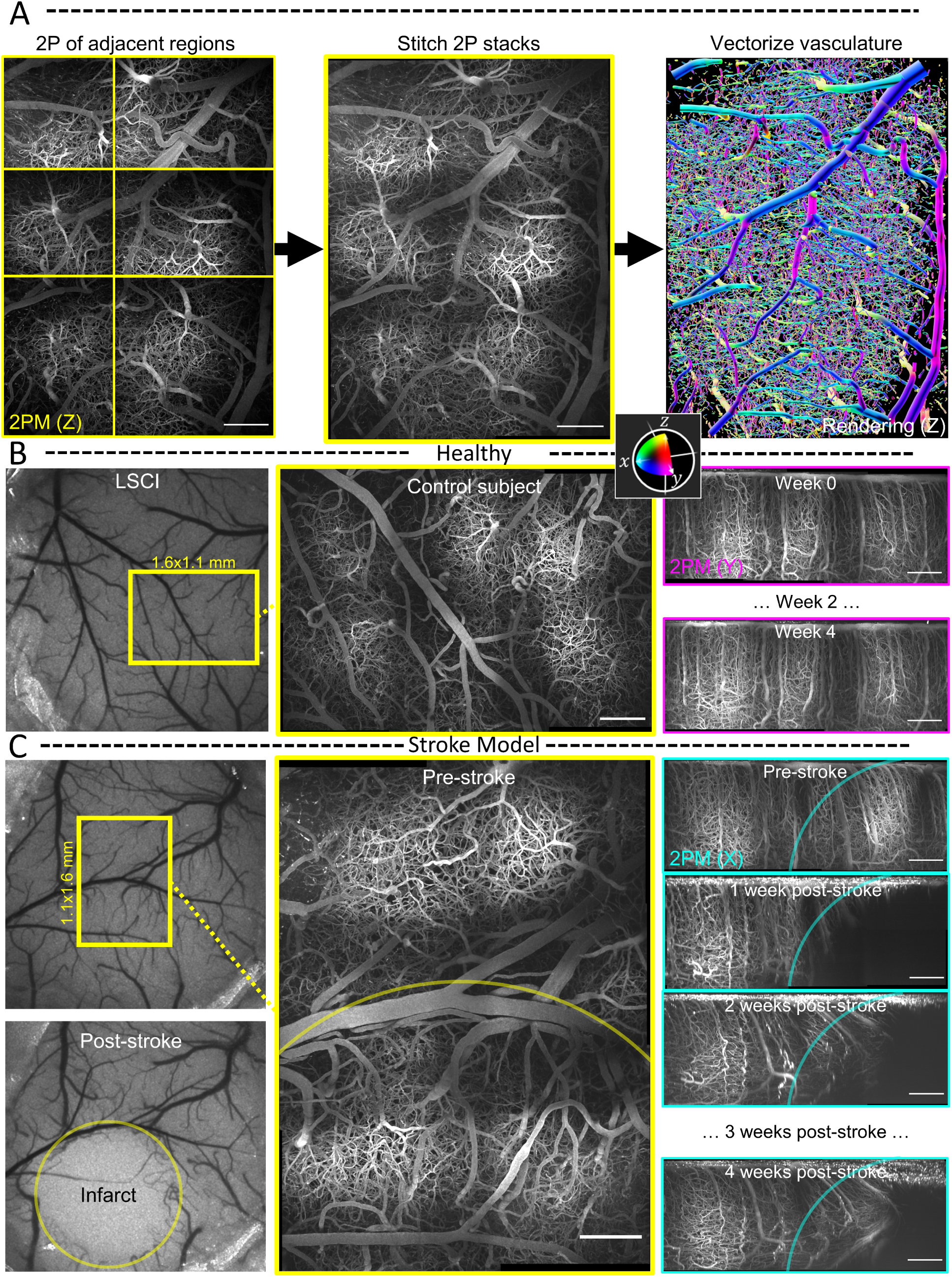
Experimental Methods: A. Mouse cortex is imaged in vivo through a cranial window using a tiling protocol to cover as large of a volume as possible in a single imaging session. The stitched image is vectorized using the SLAVV software for reconstruction, visualization, and statistical analysis, The vessel directions (as well as the borders of the 2PM images and the ROI borders) are color-coded with respect to their alignment with the imaging coordinate system: (*xyz ↔* CMY). B. LSCI is used to orient the 2PM imaging session to reproducibly image the same (1 cubic millimeter) volume longitudinally over several imaging sessions at two-week intervals and at a depth greater than 600 micrometers. Orthographic projection in the (optical) *z*-axis of a tiled image volume of a healthy control subject. Lateral orthographic projections show the longitudinal imaging reproducibility. C. The longitudinal experiment is repeated around a photothrombotic injury. The infarct appears to be contained to a (transparent yellow) circle of radius approximately 800 micrometers in the LSCI post-stroke image. The *x*-projected 2PM image reveals that a (transparent cyan) spherical ROI beneath the surface approximates the shape of the infarct.

### 2.5 Vascular Vectorization

2PM volumetric angiographs were vectorized using the Segmentation-less Automated Vascular Vectorization (SLAVV) software in MATLAB [35]. The automated vectorization outputs (mostly large vessels on the surface or penetrating) were manually (with machine learning assistance) curated into agreement with the underlying 2PM angiograph. Vascular vectorization enabled all of the statistical analyses, reconstructions, and other visualizations.

Many improvements have been made to this software since the previous publication, including improvements (A) to the manual burden of curating the automated vector output using machine learning trained on previous human curations, (B) to the design and performance of the edge extraction algorithm using a novel method inspired by the water-shed algorithm which uses the vertex objects as seed points and grows regions around each seed point while keeping track of the “best paths” home and best connections between regions while considering local vessel contrast and size, (C) to the computational efficiency of the initial linear filtering step especially for larger images and larger objects using a down-sampling protocol that limits the maximum resolution of the vessel objects in calculation.

### 2.6 Statistical Analysis of Vascular Networks

The vectorized vascular network is idealized as a collection of short contiguous cylinders “sections,” which are partitioned into the set of non-bifurcating, 1-D traces “strands,” as demonstrated in Fig. 3A. Under this idealization, anatomical statistics (vessel length, distance from ROI center, and orientation to ROI center) were calculated from the vectorized reconstructions of angiographs in MATLAB using the SLAVV software. Statistical distributions (i.e. spectra) of the vascular anatomy served to quantitatively characterize the vascular networks. Where possible, both strand- and section-based statistical (distance, orientation) distributions were generated. Strand-basis statistical (avg. radius, length, and tortuosity) distributions were generated to compare to the known previous study of microvascular network topology [23]. Section-basis statistics, which are weighted by length, were also used in the present application due to their simpler definitions, larger sample sizes, and better reproducibility.

### 2.7 Directional Analysis

Vessel directions are calculated by taking the derivative of the 3-space vessel centerline location with respect to the length coordinate along the vessel axis. Although vessel directions are color-encoded with respect to the canonical 3-space axes (Red,Green,Blue*↔ x, y, z*) only on a section basis for display purposes, for statistical purposes, the vessel orientations are summarized either on a section- or strand-basis.

### 2.8 ROI Selection

Regions of interest (ROIs) for the stroke model mice were selected to be concentric with the infarct center. Reproducible centerpoint selection was achieved by registering all volumes to a stable reference feature of the vascular anatomy (far away from the infarct). For the healthy mouse, the ROI was selected to match the geometry of the stroke mouse ROI: a sphere with center at 650 micrometer depth intersecting the rectangular imaging volume. A cylindrical ROI was used for display purposes only to help explain the directional analysis in reference to the directions of the imaging system (with the cylinder and optical axes in the *z*-direction).

### 2.9 Statistical Analysis of Longitudinal Data

The Kolmogorov-Smirnov (KS) test statistic (maximal absolute difference of CDFs) was used to summarize the differences between any two length-basis statistical distributions. Changes in vascular structure were measured by comparing longitudinal differences with respect to (A) the previous time-point and (B) the initial time-point. Baseline variations between imaging sessions of one healthy control subject, as well as between healthy subjects, were established to contextualize the response of the vascular network to a photothrombotic event.

### 2.10 Visualizations

Three-dimensional reconstructions of angiographs and two-dimensional overlays onto the maximum-intentensity-projected original 2PM images were generated using the SLAVV software in MATLAB. A custom meshing algorithm was developed and used to generate the three-dimensional vascular network reconstruction at arbitrary resolution and with a user-specified color-coding. For the purposes of this manuscript, color-coding was chosen to display the direction of each short vessel section. Direction colors are coded with respect to the imaging system (Red,Green,Blue *↔ x, y, z*) or with respect to a spherical ROI center (blue,yellow*↔*pointed toward the center,pointed tangentially to the spherical surface).

## 3 Results

### 3.1 Large-volume, 2PM in vivo angiography enables longitudinal stroke model vectorization

The imaging workflow visually described in Fig. 1A enables reproducible vascular vectorization. LSCI imaging (Fig. 1B and C, left side) was helpful in directing the delivery of a photothrombotic injury and in orienting the two-photon microscopy imaging sessions. Time-lapse volumetric images of the peri-infarct region were acquired with remarkably high quality and reproducibility to study the response of the affected neurovascular network (right side of Fig. 1B). The lateral orthographic projection time-lapse shown on the right side of Figure 1C shows the qualitative changes to the vascular network as well as to the image itself near the infarct. The image quality decreases (spatially and temporally) near the infarct with a loss of signal and an increase in background. Vessels very close (*<* 500 micrometers) to the infarct center disappear, while those nearby (*<* 1 mm) appear to be reoriented and pulled toward the infarct center.

### 3.2 Directional analysis helps visualize neuro-microvascular networks

The directions of the vessels were color-encoded In an attempt to help conceptualize the complexity of the neuro-microvascular network. The color-coding helps the image viewer appreciate the anisotropy of the vascular network, and it also facilitates vectorization performance evaluation. In Figures 1 and 2, the reconstructed color-coding highlights the vessels which are vertical (*z*-) as well as those which are aligned to the *x*- and *y*-axes of the imaging system (Fig. 2). Figure 2A shows the strand objects from an example cylindrical ROI (not used for analysis) extracted from a larger two-photon imaging volume. Figure 2B shows the agreement (within the cylindrical ROI) between the original (projected) two-photon image with the vectorized reconstruction color-coded by three-dimensional direction. The central vertical vessel featured in the vectorized reconstruction can be clearly seen in blue in all three orthographic projections.

**Fig 2.**
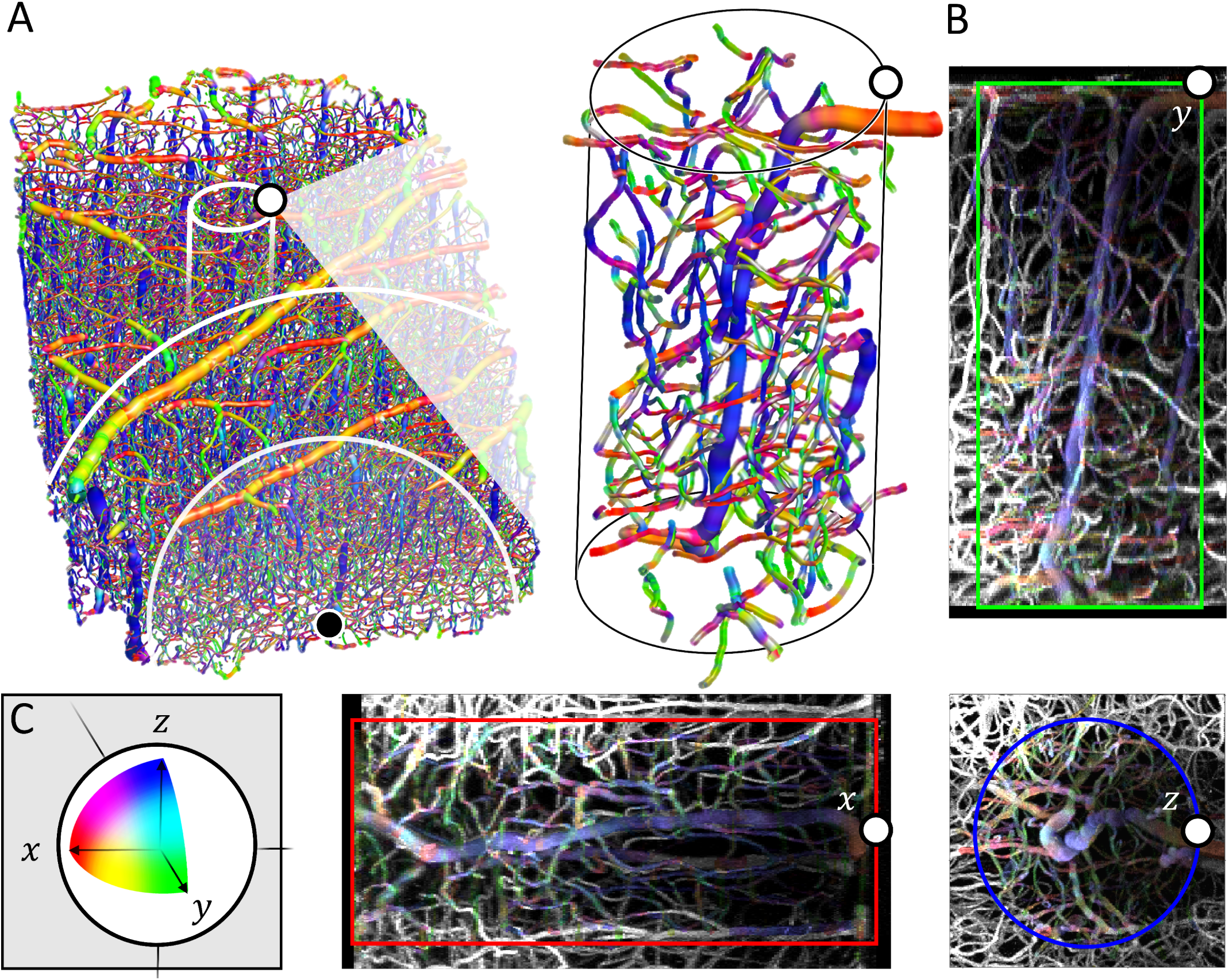
3D Directional Analysis: A. A 3D perspective rendering shows the entire capillary network captured in the rectangular 2×3 tiled image. The 1mm radius spherical ROI (500 micrometers line also shown) intersecting the near image boundary is used in the following stroke model analysis. The pop-out magnifies an example penetrating (blue) vessel and surrounding vessel strands inside of a (250 micrometer diameter, 600 micrometer height) cylindrical ROI. B. The strands from the cylindrical ROI overlay MIPs of a rectangular volume ROI of the raw 2PM image in *x*-, *y*-, and *z*-orthographic projections. C. All vessel sections in the 3D reconstructions and those belonging to strands in the cylindrical ROI in the 2D projections (as well as the ROI borders) are color-coded according to their alignment to the axes of the imaging system (*xyz ↔* RGB).

The directional information in reference to the imaging system is useful but somewhat arbitrary. In an attempt to summarize the directional information which is relevant to stroke recovery, the vessel directions were summarized by their angle in reference to the ischemic injury center. In the stroke model condition, the center of the infarct was determined by inspection of the final two-photon image, and the vessel unit vectors were projected onto the vector toward the infarct, yielding the unitless (cosine of angle) orientation statistic. The orientations of the vessels in reference to a spherical ROI are color-encoded in Figures 3C and 5A and B using a one-dimensional smooth gradient color space. Vessels which are pointed toward the infarct center appear cyan, those which are oriented perpendicular to the infarct appear yellow.

In addition to providing rich visual information, quantitative directional analysis provides a means for comparing the anatomy of two or more vascular networks. The unique signature of vessel orientation in the healthy vascular network is shown in Figure 3, where a flatline would represent an isotropic distribution of vessel directions. The two healthy mouse subjects show a similar vessel orientation signature, which becomes more obvious after normalizing the distribution by the total length to arrive at the probability distribution function (PDF shown in Fig. 4). This orientation signature is acutely unperturbed in the stroke model at one week post-photothrombosis. However, at four weeks post-stroke, the vascular network has reoriented to a new steady state which is distinct from the healthy signature. The same steady state orientation signature is seen in all three stroke model subjects at four weeks post-stroke.

**Fig 3.**
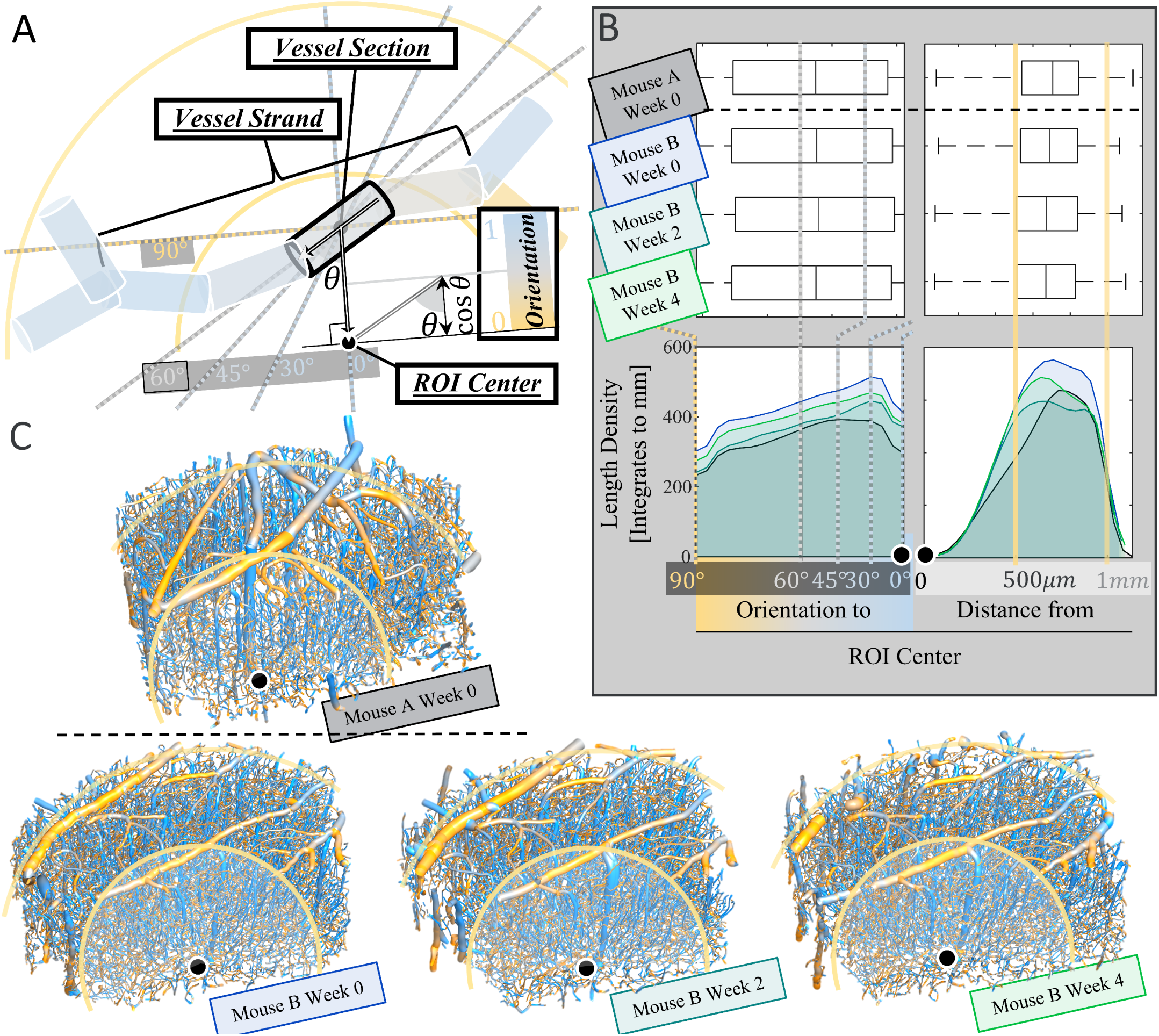
Healthy Control Snapshots: A. Legend for the directional analysis performed with respect to the spherical ROI center. Distance from the infarct center and vessel orientation are calculated for each vessel section (short vessel segment idealized as a cylinder) belonging to all strands (non-bifurcating, 1D traces) in the ROI. B. Box plots and histograms of the length-weighted distributions of orientation and distance for all of the vessel sections of all strands in the ROI. C. 3D perspective rendering of the 1 mm radius spherical control ROI matching the shape (intersected with the rectangular image volume) of the ROI used in the stroke model analysis.

**Fig 4.**
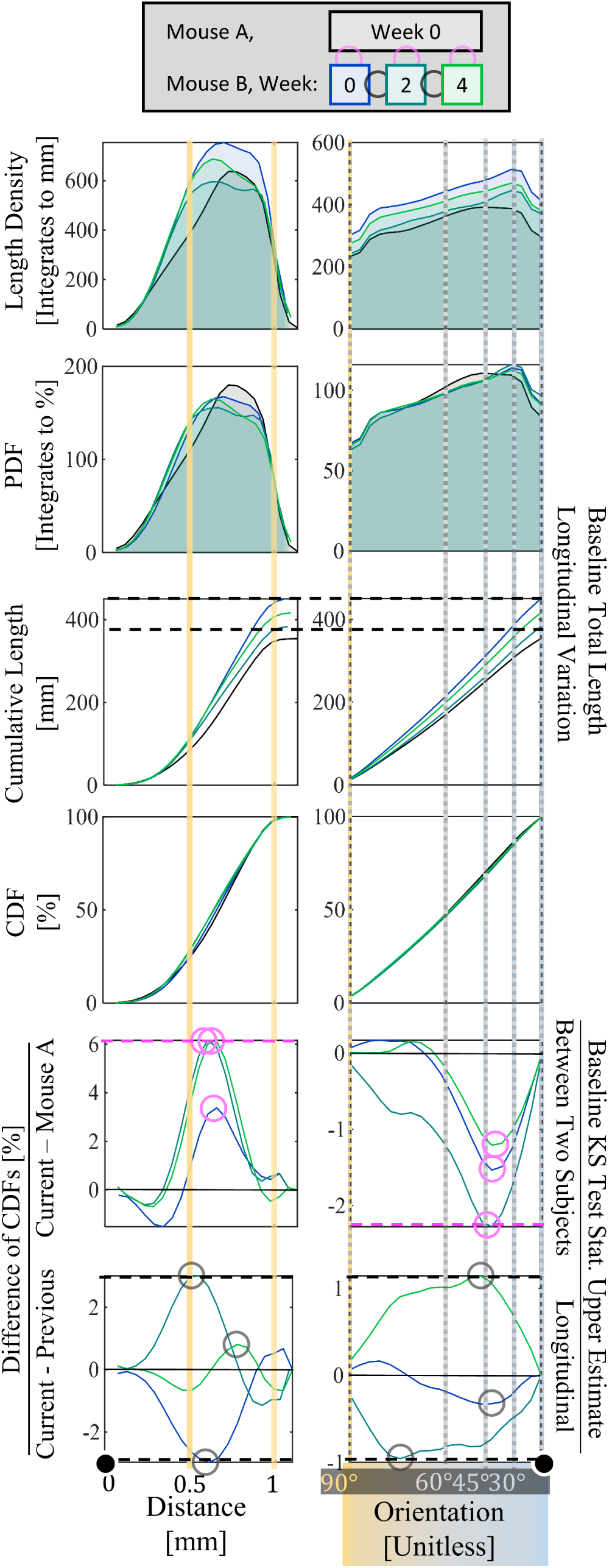
Healthy Control Statistical Analysis: From top: Four time-points from two healthy mice are compared to measure baseline statistical variations. The length densities from Figure 3 are repeated here for reference. Length densities are normalized to probability densities (PDFs) to remove differences in sample size. PDFs are integrated to cumulative distributions (CDFs), showing the variation in total length as a dotted black line which appears at the top of Figure 5C. Finally, CDFs are compared to measure the baseline variations within and between subjects. The variations between subjects A and B are larger and appear more systematic than the variations within the healthy control subject B (week 4 is assumed to be before week 0). Baseline KS test statistics are labeled with circles (and their upper estimates with dotted lines) and appear in Figure 6.

**Fig 5.**
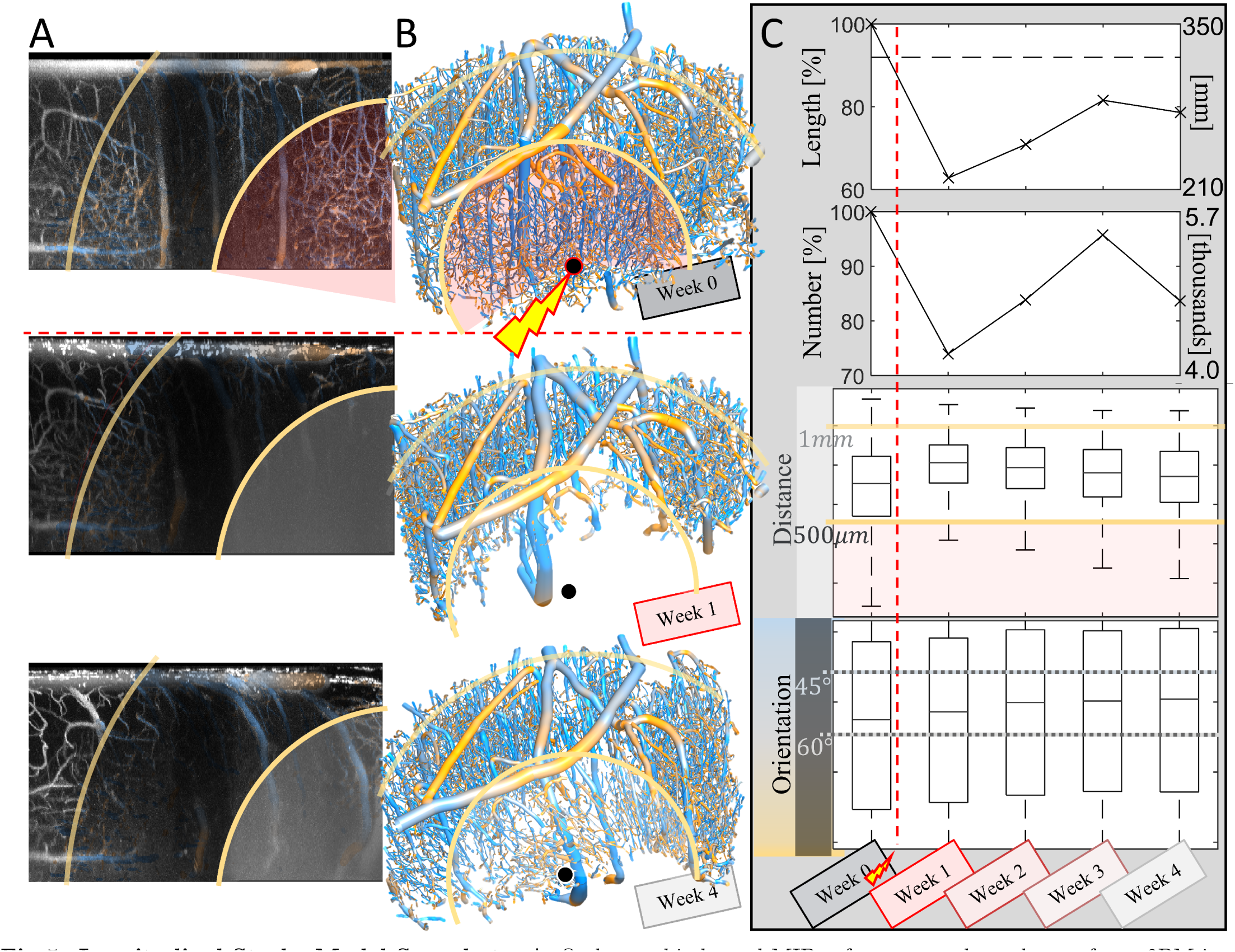
Longitudinal Stroke Model Snapshots: A. Orthographic lateral MIPs of a rectangular volume of raw 2PM image (concentric with the spherical ROI in the projection coordinate) for weeks 0, 1, and 4 post-photothrombosis. The orientation coloring for all vessel strands inside the ROI overlays the original MIP. The tissue appears to contract toward the infarct center. B. 3D perspective rendering of the strands within the spherical ROI. A smaller spherical ROI representing the core of the infarct is highlighted in red in week 0. Vessels in the core appear to be eliminated by the photothrombosis by week 1 and partially regrown by week 4. C. (From top to bottom:) total length and number of strands within the ROI and box plots of the length-weighted statistic, orientation and distance, for all vessel sections in the spherical ROI for all five imaging sessions.

### 3.3 Healthy vascular network is longitudinally stable with random statistical variations

A control mouse was imaged longitudinally to determine the healthy statistical signatures and baseline variations therefrom. A spherical ROI was selected in the control subject to match the stroke model subject ROI shape and position with reference to the larger tiled image. The vascular network was idealized as short contiguous cylinders (Fig. 3A), and summarized in terms of length-weighted statistical distributions. Figure 3B shows the signatures of distance and orientation with respect to the ROI center. Three-dimensional reconstructions with orientation color-coding in reference to the ROI center are shown in Figure 3C. The healthy control subject, Mouse B, has longitudinally stable, reproducible vectorization statistics and three-dimensional reconstructions.

The orientation and distance signatures are similar between Mouse B and the single healthy snapshot from stroke model Mouse A. Their statistical distributions can be compared in many ways: by total weight (length), median value or first moment (mean), or higher moments of the distribution (shape). Figure 4 takes a closer look at the differences between all four (one from Mouse A, three from Mouse B) healthy control snapshot distributions. To control for differences in sample size, each statistical distribution is normalized by total length to arrive at the probability distribution function (PDF), whose total area has a probability of 1 (100%).

PDFs are integrated to cumulative distribution functions (CDFs) to compare their shapes across samples and to calculate the two-sample KS test statistic, which can be conceptualized as the “distance” between statistical distributions in terms of probability (%). Longitudinal (finite) differencing of CDFs from healthy tissue shows random fluctuations in statistical distributions and represents a baseline in statistical variations (black circles and dotted lines in Figures 4 and 6B), against which the stroke model is compared. Comparison between the healthy control and the healthy time point from the stroke model subject reveals a systematic and reproducible difference signature (magenta circles and dotted lines in Figures 4 and 6B), which is larger than the random intra-subject differences, and represents an elevated baseline in statistical variations between healthy tissue samples..

**Fig 6.**
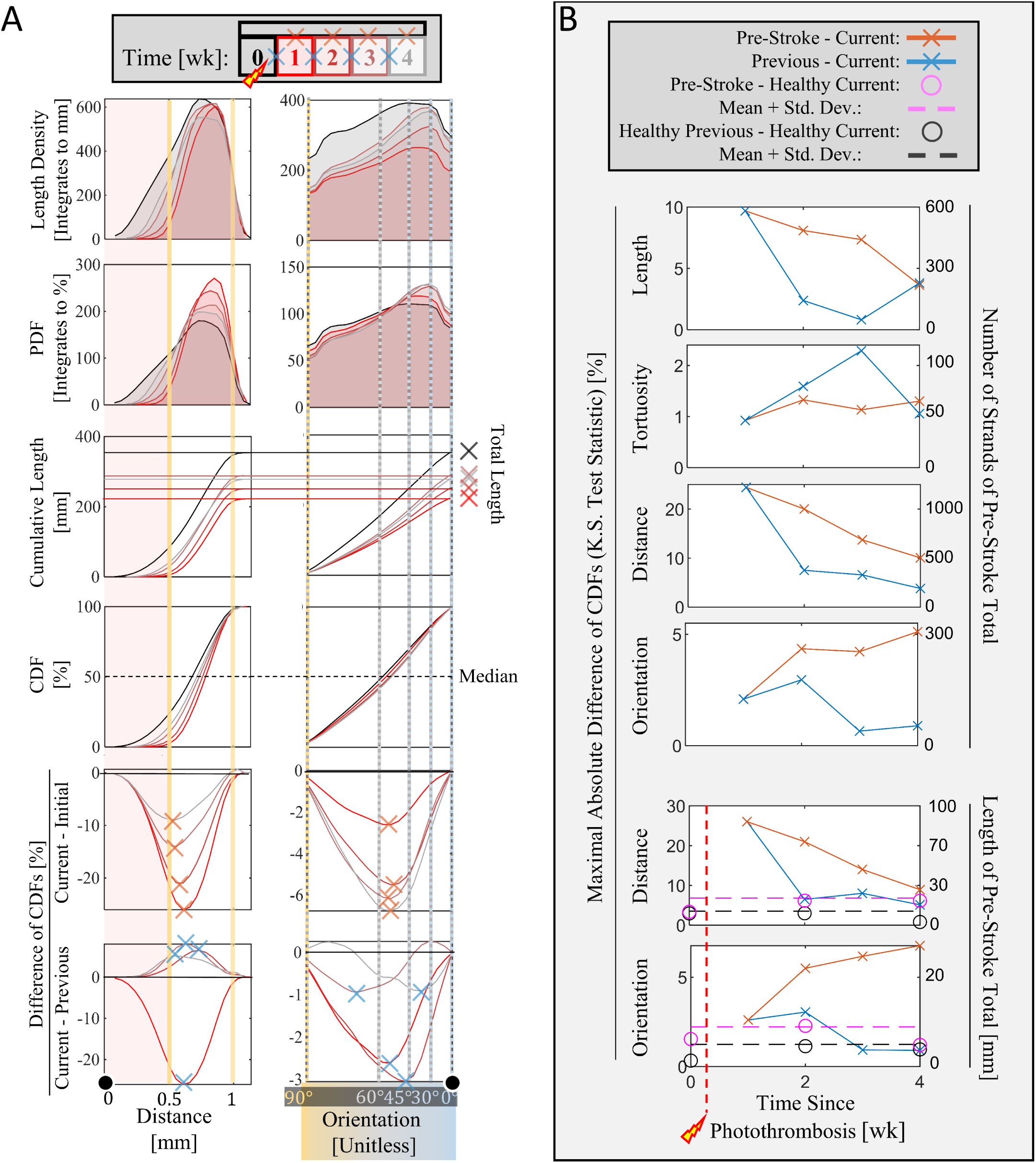
Longitudinal Stroke Model Statistical Analysis: A. Distribution snapshots of the length-weighted statistics for all timepoints in the stroke model subject (Mouse A) show the vascular response to photothrombosis. From Top: length-weighted histograms, normalized to PDF, integrated unnormalized CDF, normalized CDF, and differences of CDFs between different pairs of time-points. The extreme values of the difference of CDF plots (KS test statistics) summarize the differences between pairs of distributions. B. Quantification of pairwise differences (KS test statistic) between (top) strand-weighted and (bottom) length-weighted statistical distributions show the vascular response dynamics. The dotted lines and associated data-points represent the baseline variations within the healthy subject (Mouse B) and between healthy subjects (from Fig. 4).

### 3.4 Vascular statistical signatures are unique to pre-, intra-, and post-stroke conditions

To longitudinally track the changes in statistical distributions, five images at one week intervals from stroke model Mouse A were imaged, vectorized, and analyzed (Fig. 5). Figure 5A shows how the vascular network responds to photothrombosis. Vessels appear to be pulled toward the infarct center as the population of (blue) vessels oriented toward the infarct increases from week 1 to week 4 post-stroke. Figure 5B shows these same deformations in a three-dimensional reconstruction of the spherical ROI. The total length of vessels and total number of strands were highly correlated through this recovery process (Fig. 5C) and both take a large (30-40%) loss at week 1 and peak near (10-20% below) the baseline at the week 3 time point.

The week 1 time point is significantly different from the baseline signature in terms of distance from the infarct. The normalized orientation signature appears somewhat similar between weeks 0 and 1 (Fig. 6, PDF), however by week 4, it has trended significantly away from the baseline signature (Difference of CDFs: Current - Initial) to a new steady state (Current - Previous). Meanwhile, the distance distribution has smoothly returned to approximate the baseline signature by week 4. Therefore there are unique, measurable characteristics to each of the pre-, intra-, and post-stroke conditions.

In addition to the length-weighted statistics (distance, orientation) which are defined for short vessel sections and have a large sample size, the strand-weighted statistics (distance, orientation, radius, length, tortuosity) show more detail with a smaller sample size (Fig. 7). The distance and orientation statistics of the strand objects closely match those of the section objects, which are weighted by their length. The strand-length statistic distribution shows relatively subtle changes throughout the time-course that are highly correlated to those of the strand-distance statistic. The strand-tortuosity statistic shows a stable distribution with apparently random fluctuations around a baseline distribution, which is consistent with the healthy literature example [17, 23].

**Fig 7.**
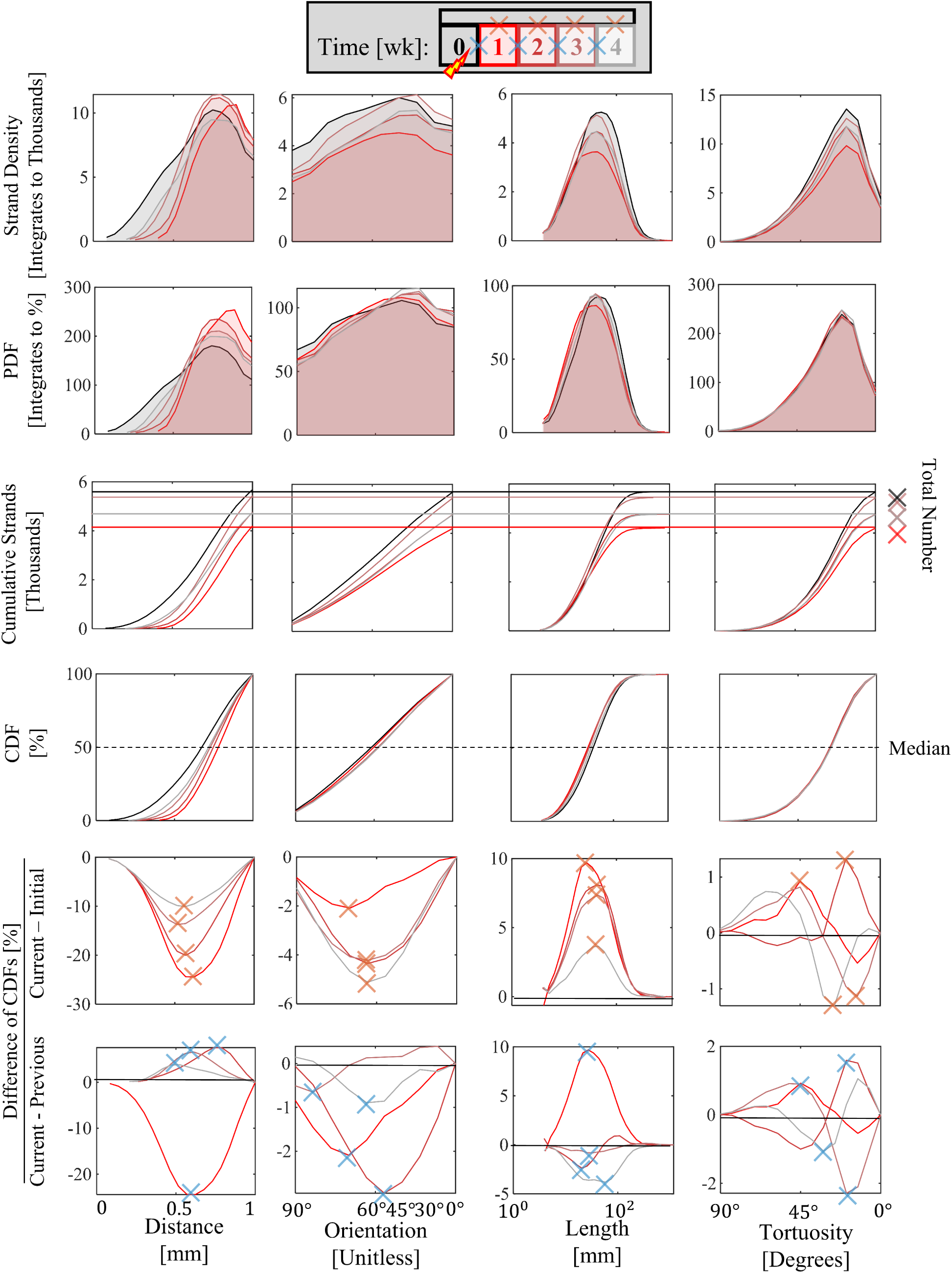
Longitudinal Stroke Model Network Analysis: Strand-weighted statistical distribution snapshots show agreement with their length-weighted version or with the healthy literature example. From Top: strand-weighted histograms are normalized to PDFs, showing increasing stability in statistics from left to right. Histograms are integrated to cumulative distributions, showing the differences in the sizes of the distributions as measured by the total number of strands. Cumulative distributions are normalized to CDFs to remove sample size differences, and then compared between pairs of time-points. The strand-length time-course shows subtle systematic variations that are correlated to those of distance. The strand-tortuosity time-course variations appear random and do not trend from the baseline.

### 3.5 Stroke model vasculature reaches a new steady state after four weeks

To determine the baseline variances within which a new steady state could be determined, the distributions from the time-course stroke model mouse A and healthy control mouse B were compared (Fig. 5, total length statistic and Fig. 6, length-weighted distance and orientation statistics). The consecutive time CDF differences (bottom of Fig. 6A) show variations within the healthy control baseline for all statistics considered at four weeks post-stroke. Furthermore, the differences in the distance and orientation statistical distributions between the pre- and four weeks post-stroke time points were larger than either the baseline inter-subject variations between the healthy pre- or between the four weeks post-stroke timepoints (Fig. 9 and 10). Therefore the vascular network has reached a new steady state at four weeks after photothrombosis.

**Fig 8.**
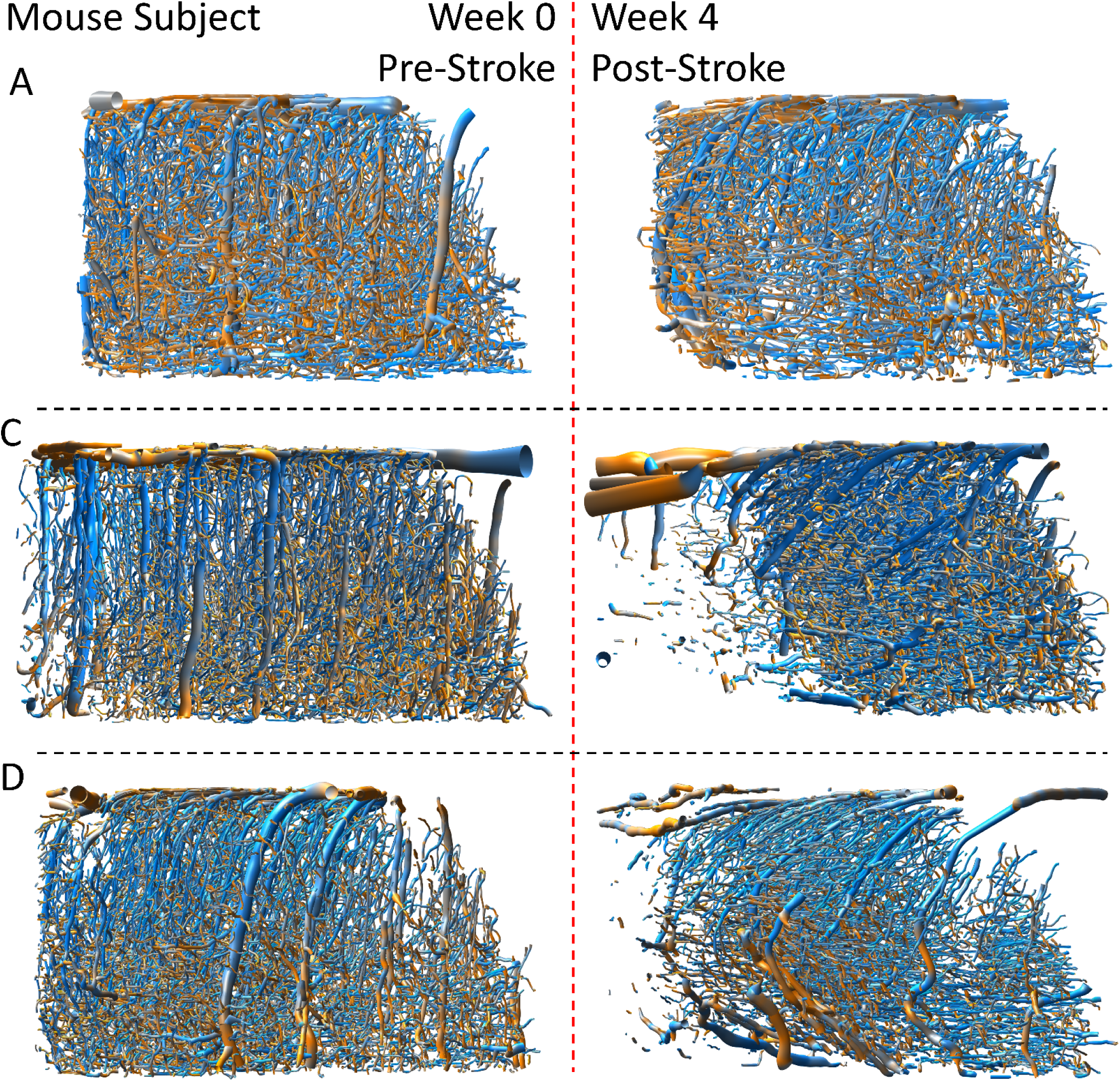
Pre- and Post-Stroke Snapshots by Subject: To compare the variation between stroke model subjects, week 0 pre- and week 4 post-stroke time-points from three mice (A, C, and D) were imaged, vectorized, and analyzed. Similar spherical ROIs were selected between subjects, concentric with the infarct center observed at week 4. Vessels appear to be removed from or reoriented and pulled toward the infarct center to different degrees between subjects.

**Fig 9.**
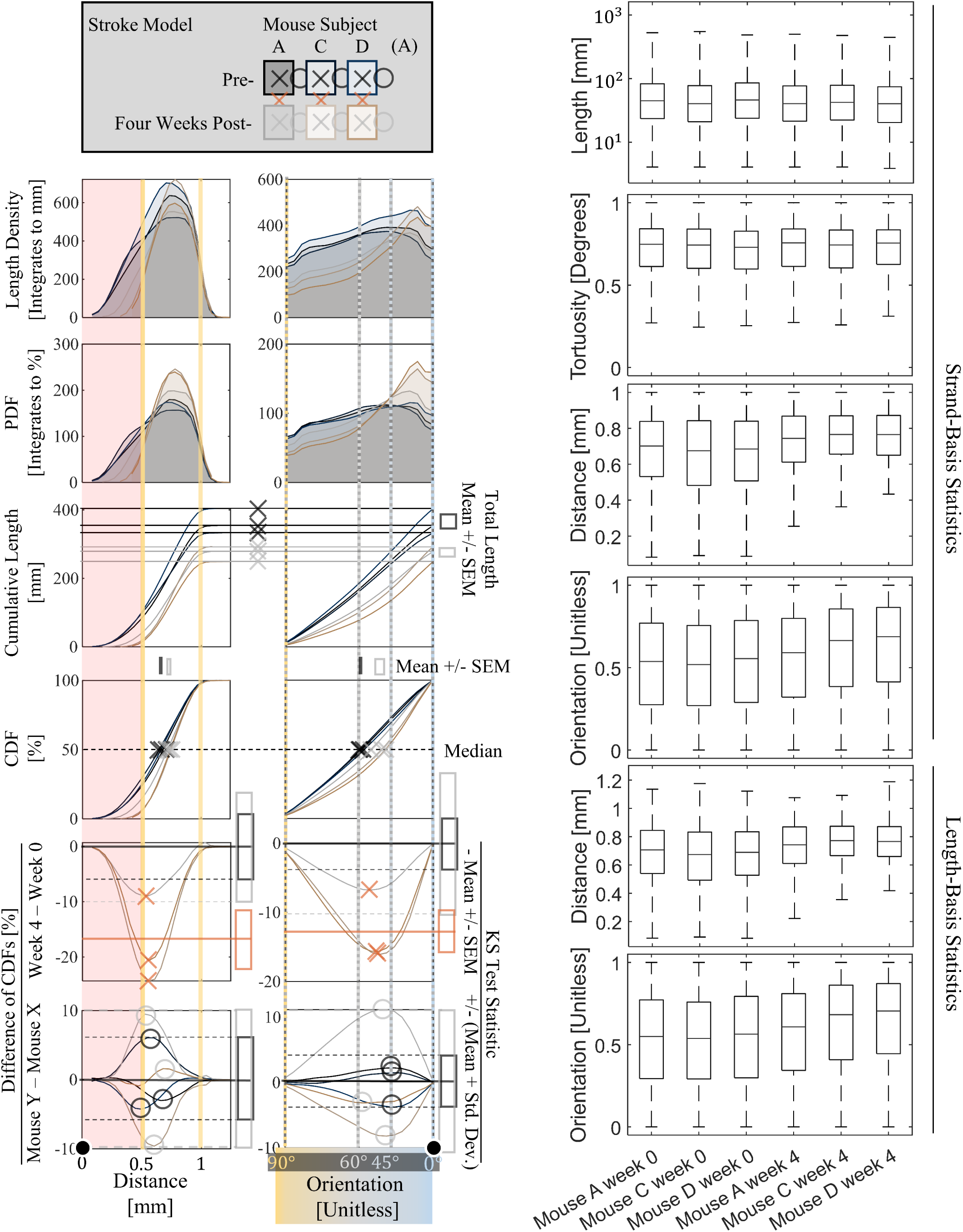
Pre- and Post-Stroke Statistical Variation: A. From top: Statistical distributions from three stroke model mice and six time-points are compared between pre- and post-injury and between subjects at either time-point. The length densities in terms of the statistics of interest (distance and orientation with respect to the infarct center) are overlaid showing a missing population near the infarct and a bias in orientation toward the infarct at four weeks post-photothrombosis. The length density is normalized to the probability density function (PDF), further highlighting the differences in the shapes of statistical distributions pre- and post-injury. The cumulative length shows the significant differences in total lengths between the pre- and post-snapshots. The cumulative length is normalized to the cumulative distribution function (CDF), revealing the significant difference between the pre- and post-injury median statistical quantities. The differences in CDFs quantifies the systematic difference in shape between the pre- and post-injury statistical distributions, revealing that the difference between pre- and post-is significantly larger than the subject-subject variation at either time-point. B. Box plots show more qualitative summary expressing the same information quantified in Panel A.

**Fig 10.**
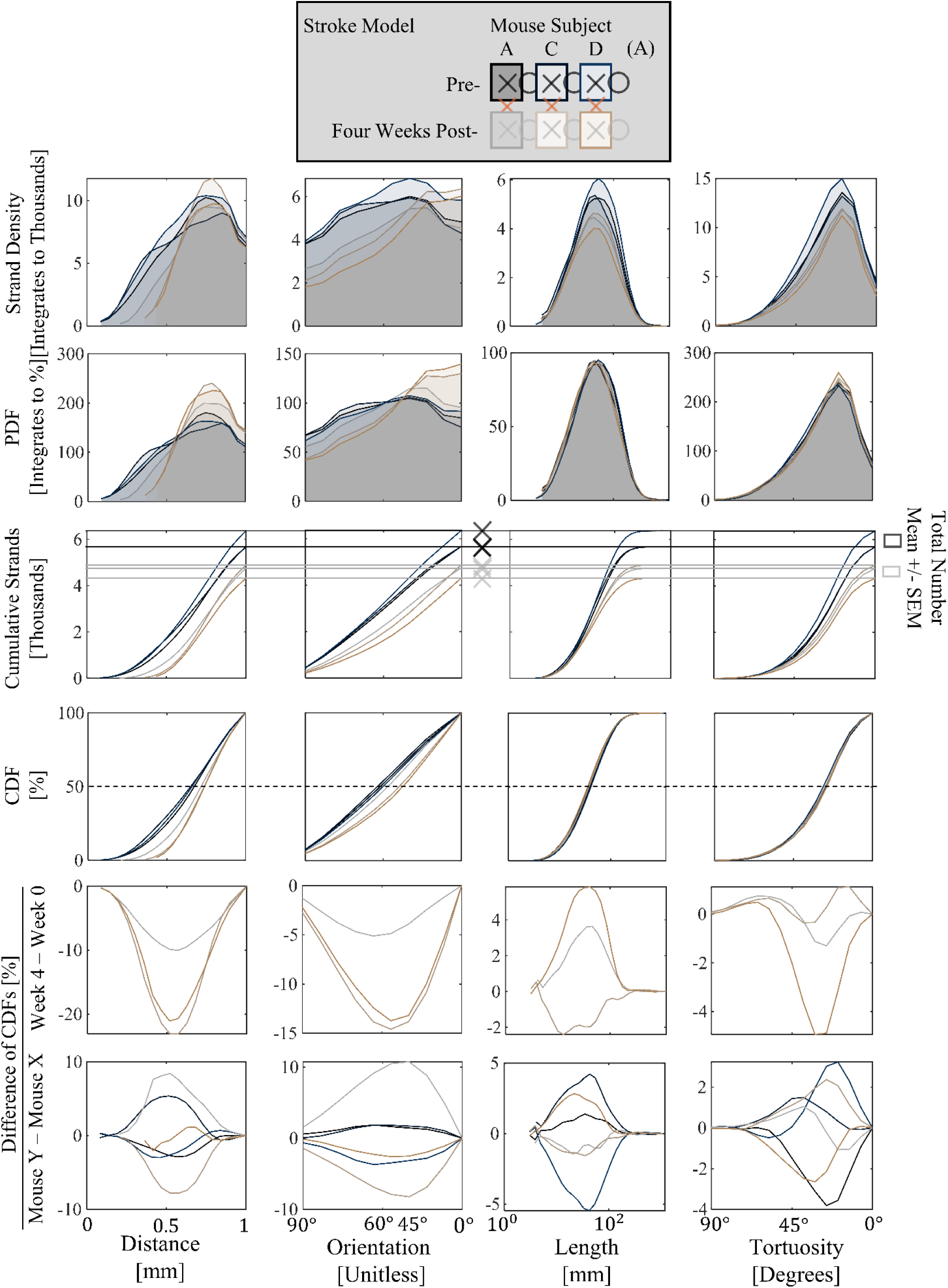
Pre- and Post-Stroke Network Variation: Strand-weighted statistical distribution snapshots show that the differences between pre- and post-stroke conditions are larger than the differences between subjects at corresponding time-points for the distance and orientation statistics, but not for the tortuosity statistic. For the length statistic, the pre-/post-stroke variations are larger than the differences between subjects at week 0 but not at week 4. From Top: strand-weighted histograms are normalized to PDFs, showing increasing stability in statistics from left to right. Histograms are integrated to cumulative distributions, showing a clear separation in the pre- and post-condition sample sizes, as measured by the total number of strands. Cumulative distributions are normalized to CDFs to remove sample size differences, and then compared between pairs of time-points to measure differences.

### 3.6 The steady state stroke model statistical signature is conserved across subjects

An additional four images at zero and four weeks post-stroke from Mice C and D were analyzed to determine the pre-/post-stroke vs. the inter-subject variation among three subjects (Fig. 8). Vectorized reconstructions with color-coded orientation with respect to the spherical ROI reveal a consistent photothrombotic event with some differences in the extent to which the surrounding tissue is affected. Vessels within about 500 micrometers from the infarct center have disappeared, while those in the surrounding sphere of 1 mm radius appear to be pulled and reoriented toward the infarct.

The statistical distribution differences between the pre- and post-stroke snapshots are larger than the differences between subjects at corresponding time-points for the (length- or strand-weighted) distance and orientation statistics but not for the strand-tortuosity statistic. The strand-length statistic has pre-/post-variations that are larger than the week 0 inter-subject differences but not larger than differences between subjects at week 4. Figures 9 and 10 show the graphical calculations of the pairwise KS test statistics to compare sample distributions and support the previous sample variation claims. The strand-radius (data not shown), -length, and -tortuosity statistics are relatively stable and consistent with the in vivo healthy mouse cortex literature examples from [17, 23]. The box plots of Figure 9B show a more qualitative summary of the above statements.

### 3.7 Stroke model neurovascular plasticity correlates with literature neuronal plasticity

According to results in a recent study of neural plasticity in a mouse stroke model citepbandet2023aberrant, the acute effects of an hypoxic injury to the neural network architecture are seen after one week, and the network has stably reorganized after four to eight weeks. The response of the neurovascular network observed in the present study correlates with this literature example, peaking in plasticity at week 1 (or 2) for the distance and strand-length (or the orientation) statistics and returning to (the new) steady state after four weeks. This correlation between neuronal and neurovascular plasticity is unsurprising in consideration of neurovascular coupling.

## 4 Discussion

### 4.1 Advantages and Disadvantages of the Experimental Tools

In this study, we employed in vivo two-photon angiography and vascular vectorization techniques to investigate the longitudinal stability of mouse cortical microvasculature and the effect of stroke. These experimental tools offer several advantages and disadvantages, which are important to consider when interpreting the results and designing future studies.

One of the major advantages of in vivo two-photon angiography is its ability to capture high-resolution images of the neuro-microvascular networks in live animals. This technique allows for the visualization of intricate anatomical features of the vasculature with a resolution of 1.5 micrometers in the *x*-*y* plane and 3 micrometers in the *z*-axis. By imaging the same region at multiple timepoints, we were able to track the changes in the vascular network over time and investigate the dynamics of stroke recovery. Additionally, the use of two-photon imaging in combination with laser speckle contrast imaging provided a means for orienting the imaging sessions and delivery of photothrombosis.

Another advantage of our study is the application of vascular vectorization, specifically the Segmentation-less Automated Vascular Vectorization (SLAVV) software, which enabled the reconstruction and quantitative comparison of the vascular anatomy across samples. This automated method significantly reduced the manual burden of analyzing the large volumes of data obtained from the imaging sessions. Moreover, improvements made to the software, such as machine learning-assisted curation of the automated vector output and computational efficiency enhancements, contributed to the accuracy and efficiency of the vectorization process. The vectorization methods have potential use cases adjacent to the intended neuro-microvasculature use case: macrovasculature in stroke [29, 20], eyes [41], ovaries [13], or in vitro [14, 15]. Additionally, the vectorized output can be exported as a standard .ply or a special .vmv file for three-dimensional rendering and analysis using the vessmorphovis citepabdellah2020interactive plugin to blender (Blender Foundation, community).

The longitudinal design of our study, with weekly imaging sessions over a period of one month, provided valuable insights into the stability and plasticity of the neuro-microvascular networks. By comparing statistical distributions of the vectorized network longitudinally and between mouse subjects, we were able to identify anatomical statistical signatures of healthy and stroke model vascular networks, as well as the dynamics of recovery. This longitudinal approach allowed us to capture temporal changes that would have been missed in endpoint studies, providing a more comprehensive understanding of cerebrovascular plasticity.

Despite these advantages, there are also limitations associated with the experimental tools used in this study. First, in vivo two-photon angiography has restrictions in terms of the lateral field of view and imaging depth. Although we were able to cover large tissue volumes exceeding 1 cubic millimeter, the imaging range is still limited compared to the entire brain. This limitation may result in a partial representation of the vasculature and could introduce sampling biases.

Another limitation is the reliance on manual curation of the automated vector output. While machine learning assistance helped to reduce the manual burden, there is still a subjective element in the curation process. Human biases and variability may influence the accuracy and reproducibility of the vectorization results. Further advancements in automated vectorization algorithms and quality control measures are needed to mitigate these limitations.

Additionally, the underlying mechanism of the vascular response to stroke, whether it is neuroplasticity, tissue deformation due to necrosis, or imaging quality artifacts remains unclear. The experimental tools used in this study provide valuable anatomical and statistical information, but they do not directly elucidate the underlying biological processes. Further investigations combining these tools with molecular and cellular techniques could provide a more comprehensive understanding of the mechanisms driving vascular remodeling and recovery after stroke.

In conclusion, the experimental tools employed in this study, including in vivo two-photon angiography and vascular vectorization, offer significant advantages in investigating the longitudinal stability of mouse cortical microvasculature and the effect of stroke. These tools enable the visualization and quantitative analysis of the intricate neuro-microvascular networks, providing valuable insights into cerebrovascular plasticity. However, limitations related to imaging range, manual curation, and the need for complementary techniques should be considered when interpreting the results and planning future studies.

### 4.2 Length-vs Strand-Basis Statistical Analysis

In our study, we performed a statistical analysis of the vascular networks based on two different approaches: length-basis and strand-basis analysis. These analyses aimed to capture different aspects of the vascular architecture and provide complementary information about the effects of stroke and longitudinal stability.

The length-basis analysis focused on measuring the total vessel length and vessel length distributions within the vascular network. This approach provides insight into the overall structural changes in the vasculature. By comparing the length distributions between healthy and stroke-affected samples, we observed significant differences in distance and orientation, indicating vascular remodeling following stroke. Specifically, we observed a decrease in vessel length and reorientation of surrounding vessels toward the infarct in the stroke-affected samples compared to the healthy controls. This finding suggests a loss or pruning of vessel segments, followed by re-growth which is consistent with previous studies on post-stroke vascular remodeling (santamaria2020remodeling). The strand-basis analysis, on the other hand, considered the individual vessel segments or strands within the network. By quantifying various parameters such as the number of strands and strand-length and -tortuosity, this analysis offers a more detailed understanding of the network’s organization. In our study, we found that the stroke-affected samples exhibited a decrease in the number of strands and an altered strand-length signature compared to the healthy controls, while the strand-tortuosity and -radius signatures were apparently unaffected. These findings suggest a disruption in the network connectivity and indicate a reorganization of the vascular architecture following stroke.

By employing both length-basis and strand-basis analyses, we gained a comprehensive view of the vascular changes associated with stroke. The length-basis analysis provided an overall measure of vascular remodeling, highlighting the loss of vessel segments, while the strand-basis analysis revealed the alterations in network connectivity and branching patterns. Together, these analyses suggest a complex restructuring of the vascular network in response to stroke.

While the length-basis and strand-basis analyses provide valuable insights, they also have certain limitations. Both approaches rely on the accuracy and completeness of the vascular vectorization process, which, as mentioned earlier, may be subject to manual curation biases. Vectorization errors would propagate to the statistical analysis with a greater effect, due to its smaller sample size and more complicated definition. The length-basis statistics have a simple definition, extremely large sample size, and are therefore somewhat robust to incomplete or inaccurate vectorization. However, the length-basis statistics do not consider network topology and are therefore lacking in the network qualities they can quantify.

In summary, the length-basis and strand-basis statistical analyses employed in our study offered distinct perspectives on the effects of stroke on the cortical microvasculature. The length-basis analysis captured the overall structural changes, while the strand-basis analysis provided insights into the network organization and connectivity. These analyses contribute to our understanding of the complex vascular remodeling processes following stroke and lay the foundation for further investigations in this field.

### 4.3 Microvascular Anatomical Statistical Analysis Interpreted

The results obtained from our study provide valuable insights into the acute microvascular changes associated with stroke, the remodeling processes, and potential implications for stroke recovery and rehabilitation. When interpreting the results, we can draw a correlation between our findings and the existing literature on vascular and neuronal plasticity in stroke models.

The pre-stroke condition is characterized by approximately uniform distributions of blood vessel length-density and orientation with reference to the (somewhat arbitrarily chosen in the case of the healthy subject) ROI center. The acute effects of the stroke are observed in the intra-stroke condition, with a significant drop in vessel length detected near the infarct center, however the orientation signature is largely unchanged. The post-stroke condition has a similar distance distribution to that of the pre-stroke condition, however, it has a different orientation signature which appears to be stable and unchanging by the end of the four weeks. These results are consistent with at least two possible explanations of neurovascular dynamics. One possibility is that the blood vessels are responding to the infarct by reorienting and growing towards the infarct. Another possibility is that the necrosed tissue is collapsing and pulling surrounding tissue toward the infarct center. This open question could possibly be answered by registering vectorized datasets and identifying and tracking vessels between time-points.

The plasticity dynamics observed in this study are consistent with a previous study which employed manual capillary tracking [53]. The observed decrease in vessel length in the stroke-affected samples compared to healthy controls, as revealed by the length-basis analysis, suggests a loss or pruning of vessel segments following stroke. This finding is consistent with the concept of vascular remodeling, which involves structural modifications in response to ischemic events [43, 55]. The reduced vessel length may reflect the retraction or elimination of vessel segments that are no longer necessary for the altered blood flow dynamics in the affected region. This remodeling process could be attributed to various factors, including reduced metabolic demands and changes in tissue perfusion patterns.

Moreover, the altered strand-length signature and decreased number of strands observed in the strand-basis analysis indicate a disruption in the network connectivity and organization following the stroke. The vascular network is a highly interconnected system that ensures efficient blood supply to the brain [38]. The observed changes in the network strand-length signature suggest a loss of complexity and compromised blood flow redistribution. The lasting alterations in network architecture observed in the orientation signature may contribute to the development of persistent functional deficits in stroke survivors, as the changes to network connectivity could impair neurovascular coupling.

Understanding the biological implications of the observed vascular changes is crucial for devising effective stroke treatment and rehabilitation strategies. The remodeling of the vasculature following stroke may have both beneficial and detrimental effects on stroke recovery [19]. On one hand, vascular remodeling can help optimize blood flow distribution and improve tissue perfusion in the affected area. On the other hand, excessive vascular remodeling may lead to chronic hypoperfusion, impairing neuronal recovery and exacerbating functional deficits.

The findings from our study highlight the need for targeted interventions that modulate vascular remodeling processes to promote optimal recovery after stroke. Impairing vascular remodeling was shown to impede blood flow re-establishment and worsen functional recovery [54]. Therapeutic approaches that promote angiogenesis, vascular maturation, and network stabilization could potentially enhance blood flow restoration and support neuronal regeneration in the post-stroke period [37]. Additionally, interventions aimed at enhancing collateral vessel growth and improving network connectivity may facilitate functional recovery by providing alternative routes for blood supply.

Furthermore, the insights gained from our statistical analyses can inform the development of computational models and simulations of the post-stroke vascular network response [24]. These models can help investigate the complex interactions between vascular remodeling, tissue reorganization, and functional recovery. By integrating experimental data with computational models, we can gain a deeper understanding of the underlying mechanisms driving the observed vascular changes and their impact on stroke outcomes.

Results from our study on neurovascular plasticity after stroke correlate with existing literature on neuronal plasticity in stroke models [42, 2] and hypoperfusion [11]. Similar to previous findings, we observed acute effects of a hypoxic injury after one week, with neurovascular plasticity peaking at one week by the distance and strand-length statistics and two weeks by orientation. The neurovascular network reached a distinct stable state with a distinct orientation signature after four weeks, mirroring the stabilization seen in the neural network.

This correlation between neuronal and neurovascular plasticity is expected due to neurovascular coupling, which ensures the coordination of neuronal activity and blood flow regulation [51]. Our findings reinforce the idea that stroke-induced remodeling involves both the neural and vascular components of the brain. Understanding the interplay between neuronal and neurovascular plasticity is crucial for developing effective therapeutic strategies for stroke recovery.

In conclusion, the results interpreted from our study shed light on the vascular alterations associated with stroke and provide a foundation for future investigations in stroke recovery and rehabilitation. The observed changes in vessel length density and network architecture offer valuable insights into the vascular remodeling processes following stroke. These findings have important implications for developing targeted interventions and computational models that aim to optimize stroke recovery and improve patient outcomes. Our results emphasize the dynamic nature of the neurovascular network after stroke and highlight the importance of considering both neural and vascular remodeling in response to stroke. Future research should delve into the molecular and cellular mechanisms underlying these changes and explore interventions to modulate neurovascular remodeling.

### 4.4 Possible Applications of Novel Experimental Tools

The novel experimental tools utilized in this study offer several potential applications in the field of neurovascular research. By combining advanced imaging techniques with statistical analysis, we gain valuable insights into the complex interactions between neural and vascular networks.

1. Stroke Rehabilitation: Understanding the dynamics of neurovascular remodeling can aid in the development of innovative rehabilitation strategies [1, 44]. By assessing the changes in neurovascular architecture over time, these tools can help identify critical time windows for interventions and assess the effectiveness of different rehabilitation approaches.
2. Drug Development: The ability to quantify neurovascular plasticity provides a valuable tool for assessing the efficacy of drugs targeting stroke-induced remodeling [37]. These tools can help researchers evaluate the impact of pharmaceutical interventions on both neural and vascular parameters, guiding the development of new therapeutics.
3. Neurodegenerative Diseases: Neurovascular dysfunction is a common feature in many neurodegenerative diseases, such as Alzheimer’s and Parkinson’s [56]. The application of these experimental tools can provide insights into the alterations of neurovascular coupling in these conditions, aiding in the development of early diagnostic markers and potential therapeutic targets.
4. Brain Development and Aging: The investigation of neurovascular plasticity using these tools can shed light on the normal aging process and age-related changes in the brain [32]. By comparing neurovascular remodeling patterns in different age groups, researchers can gain a better understanding of the mechanisms underlying cognitive decline and develop interventions to promote healthy brain aging.
5. Brain-Computer Interfaces (BCIs): BCIs are emerging technologies that enable direct communication between the brain and external devices [10]. The knowledge gained from studying neurovascular plasticity can enhance the design and implementation of BCIs, improving their accuracy and reliability by considering the neurovascular changes induced by brain-computer interactions.
6. Personalized Medicine: Each individual may exhibit unique patterns of neurovascular plasticity [33]. These experimental tools can contribute to personalized medicine approaches by allowing for the assessment and monitoring of an individual’s specific neurovascular remodeling in response to stroke or other neurological conditions. This information can guide personalized treatment strategies and optimize patient outcomes. The novel experimental tools employed in this study have promising applications in stroke rehabilitation, drug development, neurodegenerative diseases, brain development, brain-computer interfaces, and personalized medicine. By advancing our understanding of neurovascular plasticity, these tools have the potential to revolutionize clinical approaches and improve outcomes in various neurological conditions.

## 5 Conclusion

This research article has explored the dynamics of cerebrovascular plasticity by tracking anatomical and topological statistical distributions in a stroke model using a high-throughput and high-performance in vivo two-photon angiography and vascular vectorization method. The findings shed light on the intricate interactions between neuro-microvascular networks and their remodeling following a stroke event. The results demonstrate that neurovascular plasticity occurs in response to stroke-induced injury, with significant changes observed in both the spatial organization and length distribution of the vascular network within one week of the ischemic injury.

The application of advanced imaging techniques and statistical analysis has allowed for a detailed characterization of the neurovascular remodeling process. The findings highlight the acute effects of hypoxic injury, with changes in the vascular network architecture becoming apparent within one week. Moreover, the stability and reorganization of the network were observed to stabilize by four weeks, but with enduring consequences. These temporal patterns align with existing literature on neuronal plasticity, further emphasizing the correlation between neuronal and neurovascular remodeling in stroke models.

The development and utilization of a novel longitudinal experimental protocol, including in vivo two-photon angiography and vascular vectorization, has provided valuable insights into the complex dynamics of neurovascular plasticity. The tools demonstrated and explained here offer potential solutions to problems in various areas of research and clinical practice. In stroke rehabilitation, understanding the time windows of neurovascular plasticity can guide the development of effective rehabilitation strategies. Additionally, the assessment of neurovascular remodeling can aid in evaluating the efficacy of drugs targeting stroke-induced changes and exploring potential therapeutic interventions.

Furthermore, the experimental methods have applications beyond stroke research. Neurodegenerative diseases, brain development, aging, brain-computer interfaces, and personalized medicine can all benefit from studying neurovascular plasticity. The characterization of neurovascular remodeling patterns in these contexts can facilitate early diagnosis, identify therapeutic targets, improve brain-computer interface designs, and enable personalized treatment strategies.

In summary, this research article has provided significant insights into the statistical details of cerebrovascular plasticity and its correlation with neuronal plasticity in a stroke model. The utilization of in vivo two-photon angiography and vascular vectorization has advanced our understanding of neurovascular remodeling, offering potential applications in stroke rehabilitation, drug development, neurodegenerative diseases, brain development, brain-computer interfaces, and personalized medicine. The findings contribute to a broader knowledge of neurovascular interactions and pave the way for future studies aiming to unravel the complex mechanisms underlying neurovascular plasticity in various neurological conditions.

## Supporting information

**S1 Appendix. SLAVV software source code (MATLAB)** https://github.com/UTFOIL/Vectorization-Public

## Acknowledgments

This work was supported by the NIH. Aaron Woods and Alankrit Tomar helped with the manual curation burden.

